# Decoding representations of discriminatory and hedonic information during appetitive and aversive touch

**DOI:** 10.1101/2020.09.24.310383

**Authors:** James H. Kryklywy, Mana R. Ehlers, Andre O. Beukers, Sarah R. Moore, Rebecca M. Todd, Adam K. Anderson

## Abstract

Emotion is typically understood to be an internal subjective experience originating in the brain. Yet in the somatosensory system hedonic information is coded by mechanoreceptors at the point of sensory contact before it reaches the central nervous system. It remains unknown, however, how these distinct peripheral channels for tactile hedonic information contribute to representations of interoceptive states relative to exteroceptive experience. In this fMRI study we applied representational similarity analyses with pattern component modeling, a technique that deconstructs representational states into a weighted set of distinct predefined constructs, to dissociate how discriminatory vs. hedonic tactile information, carried by A- and C-/CT-fibers respectively, contributes to population code representations in the human brain. Results demonstrated that information about appetitive and aversive tactile sensation is represented separately from non-hedonic tactile information across cortical structures. Specifically, although hedonic touch originates as a peripheral signal, labeled at the point of contact, representations in somatosensory cortices are guided by experiences of non-hedonic touch, By contrast, representations in regions associated with interoception and affect encode signals of hedonic touch. This provides evidence of complex tactile encoding that involves both external-exteroceptive and internal-interoceptive dimensions. Importantly, hedonic touch contributes to representations of internal state as well as those of externally generated stimulation.

## Introduction

Sensory experiences, such as the embrace of a loved one or the pain of a stubbed toe, can be broken down into two central components: discrimination of the sensory information and the associated hedonic response. The information processed by sensory systems is typically viewed as objective, forming representations of a tangible external environment. In contrast, hedonic appraisal is a subjective centrally mediated process (Rolls, 2019; Todd et al., 2020) for estimation and comparison of affective value. However, there is evidence that hedonic information (good vs bad) is coded by the peripheral afferents of the somatosensory system prior to any cortical processing (Iggo, 1959, 1960; Vallbo et al., 1999) suggesting that aspects of tactile sensation are valenced from the point of contact (Miskovic & Anderson, 2018). It has been proposed that these signals of peripherally identified hedonic information contribute to internal representations of homeostatic threat and social safety (Craig, 2011, 2015), rather than to traditionally somatosensory representations (i.e., a physical object contiguous to our person), yet this notion remains largely untested. If found to be true, this would indicate a unique dual role of our proximal senses consistent with Sherrington’s classic distinction between *exteroception*, or sensation of an object in the external environment, and *interoception*, which entails sensing the body itself as object (Sherrington, 1906). The current work aims to examine these hypothesized neural dissociations between exteroception and interoception in the experience of touch. We assessed whether affective qualities of aversive pressure and caress are represented distinctly from discriminative information. Such a dissociation would suggest anatomically distinct tactile signalling pathways for hedonic tactile information, reflecting decentralized affective processing (Kryklywy et al., 2020) in brain regions distinct from exteroceptive somatosensory cortices.

The somatosensory system contains multiple functional subsystems, with specific peripheral nerves serving as labeled lines for information traveling into the central nervous system (McGlone & Reilly, 2010; McGlone et al., 2014). Fast large-diameter myelinated afferent fibers (A-fibers) support sensory discrimination. These fibers predominantly convey information about the timing and location of cutaneous sensory stimulation (McGlone & Reilly, 2010), with some fibers specialized for nociception (Nagi et al., 2019). Additional small-diameter unmyelinated afferent pathways (C-Fibers; Qiu et al., 2006) support the hedonic response to touch, conveying information about affective aspects of aversive touch and nociception. At the cortical level, primary somatosensory cortex (S1) is the dominant entry point for information carried along myelinated cutaneous pathways. There is evidence that integration of hedonic information into discriminatory touch representations in these early sensory structures occurs through centrally-mediated appraisal of pain and pleasure (Bushnell et al., 1999; Gazzola et al., 2012). This observation is consistent with conventional views positing that modulation of sensory information by emotion is a centrally mediated process that relies on re-entrant projections from higher order structures assessing hedonic value (e.g., prefrontal cortices [PFC], insula, amygdala) to the sensory cortices (Pessoa & Adolphs, 2010; Rolls, 2019).

Yet, evidence for peripheral labeling of affective information suggests that not all affective modulation of sensory signals is the result of central feedback (Qiu et al., 2006). Considerable evidence exists to support a neural bases of pain-coding in the periphery (McGlone & Reilly, 2010; Nagi et al., 2019). Anatomical projection studies in primates indicate that information carried along unmyelinated C-fiber pathways does not project to the entirety of S1, as observed for A-fibers. Rather, it projects to an anterior region of S1 (insula-adjacent area 3a; Vierck et al., 2013; Whitsel et al., 2009), with additional direct projections to the insula, anterior cingulate cortex (ACC), and PFC (Baumgartner et al., 2006; Qiu et al., 2006). Similarly, recently identified pathways – C-tactile (CT) fibers – have been shown to carry information about caress or pleasant touch (Loken et al., 2009; Marshall et al., 2019). These fibers originate from mechanoreceptors located in hairy skin rather than the glabrous (i.e., hairless) skin of the palms, where previous research has focused (Marshall et al., 2019; McGlone & Reilly, 2010; McGlone et al., 2014). These CT-fiber afferents respond preferentially to touch that is subjectively perceived as pleasant caress (Croy et al., 2016; Loken et al., 2009; Olausson et al., 2002). Thus, in the cutaneous system there may be distinct parallel representations for tactile stimulation beginning from the point of contact, carried through hedonic labeled lines, that independently inform the experience of hedonic value.

Previous functional Magnetic Resonance Imaging (fMRI) studies examining neural substrates of affective-tactile processing have investigated either C- or CT-fiber pathways, but not both. The independent examination of C- and CT-fibers does not allow for the dissociation between these two distinct systems and is unable to discriminate valence-specific hedonic information from general tactile salience and arousal. Moreover, these studies have also relied predominantly on univariate statistical approaches (Loken et al., 2009; McGlone et al., 2014; Olausson et al., 2002). Univariate approaches have limited ability to discriminate specific information that is represented within a region, particularly perceptual and hedonic information represented by different sensory systems (Chikazoe et al., 2014; Todd et al., 2020). By contrast, multivariate analyses, including representational similarity analyses (Kriegeskorte et al., 2008) allow the examination of population-based neuronal coding in a multidimensional representational space. When further analysed through pattern component modelling, this representational space can be decomposed into weighted sub-components of experience (Diedrichsen et al., 2018), thus representing neural activity as an integration of multiple heterogeneous sets of overlapping neural representations. In the present study, we implemented an innovative analytic approach that uses theory-guided components to perform *pattern component modelling* (PCM; Diedrichsen et al., 2018; Kriegeskorte & Kievit, 2013). This PCM derivative (J.H. Kryklywy et al., 2021) works by fitting multiple theoretical similarity matrices characterizing perfect neural representation of information vectors, called here *information pattern components* (IPCs), to observed representational patterns extracted from a series of predefined regions of interest (ROIs).

In the present study, functional neuroimaging data was collected while participants received aversive pressure on the thumb or appetitive caress stimulation on the forearm (Figure 1A) and viewed images of faces with neutral expressions. RSA conducted on the BOLD signal identified similarity/dissimilarity between tactile conditions that not only represent distinct discriminatory patterns but also putatively stimulate distinct fiber pathways for aversive and appetitive tactile experience (Figure 1B). IPCs were created for task-relevant information constructs and included specific aspects of tactile and emotional experience. Bayesian information criterion (BIC) analyses were then performed to characterize the combination of IPCs that best predicted observed similarity patterns of neural activity in each ROI. This allowed us to identify and weigh dissociable representations of discriminative and hedonic tactile signals, revealing potential C-fiber and CT-fiber pathways projection.

**Figure 1.**
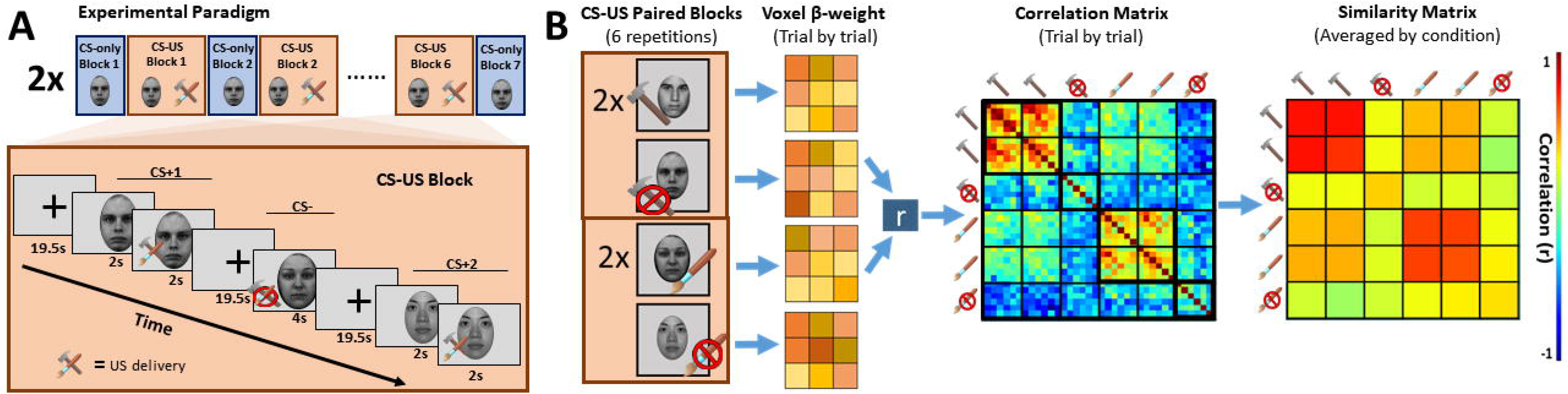
*A) Experimental time course*. Participants completed tactile visual conditioning tasks. Only data collected during CS-US paired blocked will be presented. *B) Representational similarity analyses (RSA)*. RSA was conducted correlating all experimental trials independently. Resultant Pearson correlation coefficients were averaged across conditions (removing autocorrelation) to create a 6 × 6 condition similarity matrix comparing all conditions of interest.

We predicted that representation of the hedonic components of the tactile stimulation in frontotemporal cortices, including vmPFC, ACC, and insula would be distinct from representations in primary somatosensory cortices. This would demonstrate the dual coding of somatosensation and indicate that tactile afferents coding appetitive and aversive touch are coded as internal states beyond their representation as external sensory events. We expected that dissociable representational patterns for appetitive vs. aversive tactile stimulation would be identified in the insula and vmPFC, as these regions may receive direct unprocessed information from hedonic-labeled lines. Patterns observed in the ACC were predicted to be most heavily weighted towards representation of aversive touch, consistent with this region’s preferential activation in response to modality-general pain.

## Results

To decode the neural instantiation of discriminatory and hedonic touch, we employed representational similarity analyses (RSA) with pattern component modelling (PCM) on functional neuromaging data collected during affective tactile manipulation. Three distinct tactile experiences were induced in the scanner over the course of two experimental runs. In one run, participants were presented with six repetitions of two faces paired with aversive pressure, and one face left unpaired. In the other run, participants were presented with six repetitions of two new faces paired with appetitive caress, and a third face paired left unpaired (Figure 1). Thus, there were six distinct conditions across the full paradigm (2 X aversive pressure, 2 X appetitive caress, and 2 conditions lacking tactile manipulation [once alongside each hedonic set]). RSA conducted on these data determined the degree of similarity between the voxel-by-voxel activation patterns for each condition within nine individual regions of interest (ROIs; see Table 1). We defined ROIs based on two distinct categories: *Exteroceptive* ROIs included primary somatosensory cortex, secondary somatosensory cortex, primary visual cortex and ventral visual structures while *interoceptive* ROIs included amygdalae, ventromedial prefrontal cortex, anterior cingulate cortex, and both the anterior and posterior divisions or insular cortex. Subsequent PCM iteratively fit pre-defined pattern components representing specific constructs to the observed similarity patterns in each ROI using Bayesian model fitting (Bayesian Information Criterion; BIC). Here we employed a greedy best-first search algorithm to determine the nature of information represented in a given region, as defined by the contributing pattern components, and their representational strengths. Results presented here are generated from a Monte-Carlo cross validation procedure (1000 iterations) conducted on a random sample of 60 participants (RS = 60), with identified IPCs, and their representational weights then fitted validated against the remaining seven participants (the ‘hold-out’; HO = 7). For additional detailed results about model search paths and component weighting from full sample analyses (HO = 0; Figures 3 and 4), see Supplemental Tables ST1, ST2 and ST3.

### IPC identification and weighting

To deconstruct the observed representational similarities into defined independent pattern components (IPCs), thirteen IPCs were constructed representing ideal categories of task-relevant information for use in a novel form of Pattern Component Modeling (Table 1; see Supplementary Material S1 for an extended discussion). As an example of the differences between these models, consider ‘Specific touch [ST],’ and ‘Touch valence [TV]’ (Table 2). Significant fit of ‘Specific touch’, an IPC modeling discriminative tactile information, would indicate that a region displayed a unique pattern of BOLD activation across voxels for each tactile experience encountered (i.e., appetitive touch, aversive touch and the absence of both). By contrast, significant fit of ‘Touch valence’, an IPC modeling integrated hedonic information, would involve high representational similarity within appetitive and aversive trials, but dissimilarity between these two conditions, specifically placing the tactile valence on an opposing linear spectrum (Chikazoe et al., 2014).

To determine the IPC combinations that best explained the observed correlations in the data for each ROI, PCM was conducted though Bayesian Information Criterion (BIC) analyses and an uninformed *greedy best-first search* (*GBFS*) algorithm. To ensure that regression fits were not a product of overfitting, these analyses were performed as a Monte-Carlo cross validation. Specific outputs of interest included the proportion of cross-validation iterations (i = 1000) in which an IPC was identified as a contributing component of the experimental data in the random samples, the representational weight of those components identified at a rate significantly greater than chance, and the model fit of the reconstructed RS components to the holdout (Figure 2; for complete summary of the cross-validation results, see Table 3). The proportion of iterations for IPC identification was compared to chance identification for each region of interest (ROIS, i.e., # of IPCs identified / total # of IPCs). For each iteration of the MCCV procedure, the total number of paths required for a given search is defined as the n-path.

**Figure 2.**
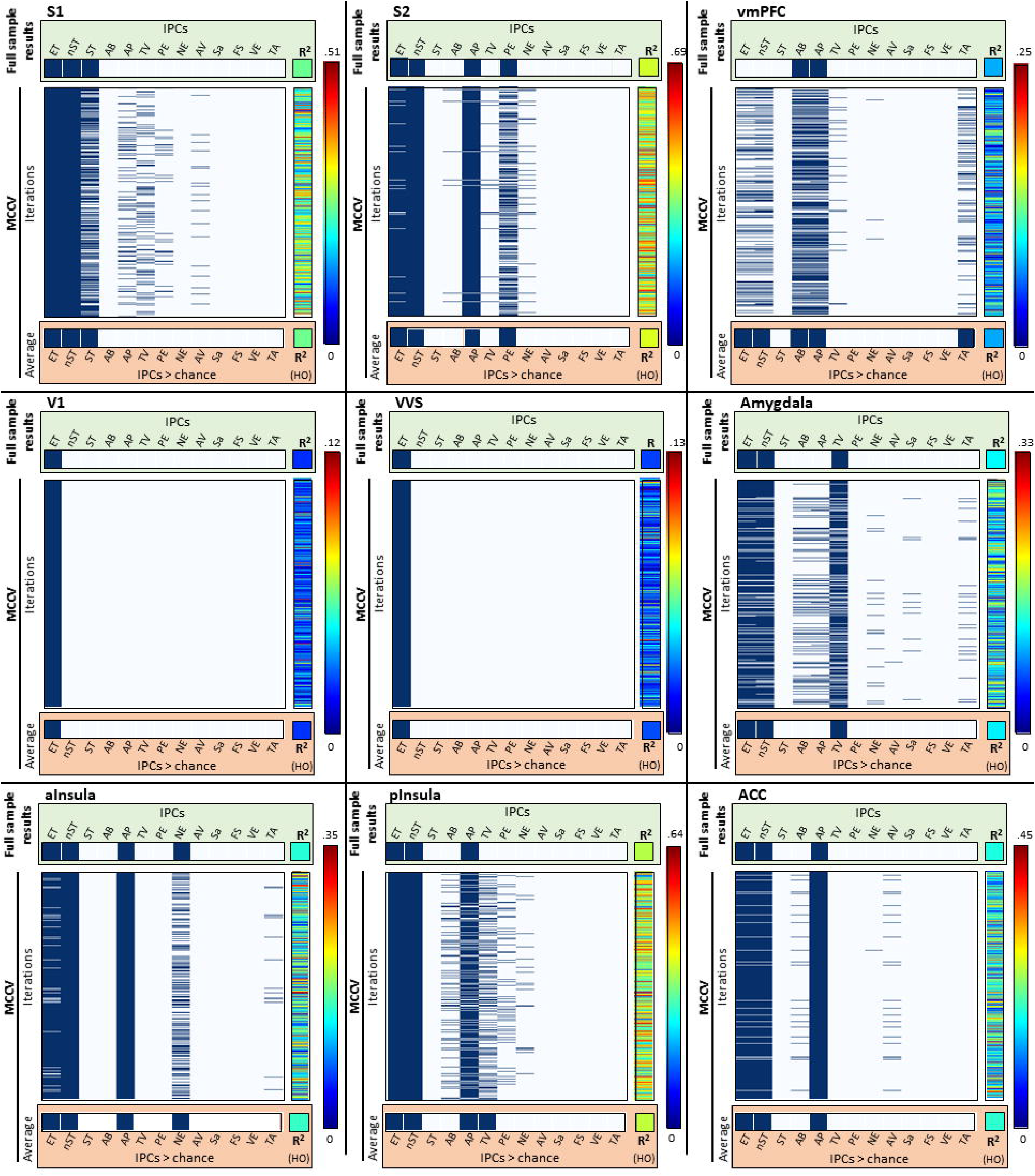
*Cross-validation across brains*. A 1000 iteration monte-carlo cross validation determined 1) that identified IPCs from the whole sample data (n = 67) were reliably identified when the procedure was replicated on subsets of the sample (n = 60) and 2) that reconstructed data generated through IPC identification and weighting accurately predicted activational similarity pattern in the held-out participants (n = 7). Results from each MCCV iteration are represented as a row of data, with the identified IPC noted and the dark blue, and the fit to the HO shown in the center-right column for each ROI. Data summaries collapsed across all MCCV iterations in shown in the red box for each ROI.

#### Exteroceptive regions of interest

##### Primary Somatosensory Cortex (S1)

In S1, a component modeling non-specific aspects of tactile experience (nST) was identified as the strongest individual predictor of S1 representational patterns in the random samples (β_nST_ = 0.129). Additional components identified at a rate significantly greater than chance modeled experimental task and non-hedonic tactile experience (β_ET_ = 0.055 and β_ST_ = 0.025 respectively). In the held-out sample, components identified in the random sample explained 24.8 % of the variance (R^2^ = .248, *F* _(1,145)_ = 52.13, *p* < .001). This pattern indicates a representation of distinct discriminative tactile experiences rather than hedonic value in S1 (Figure 3A).

**Figure 3.**
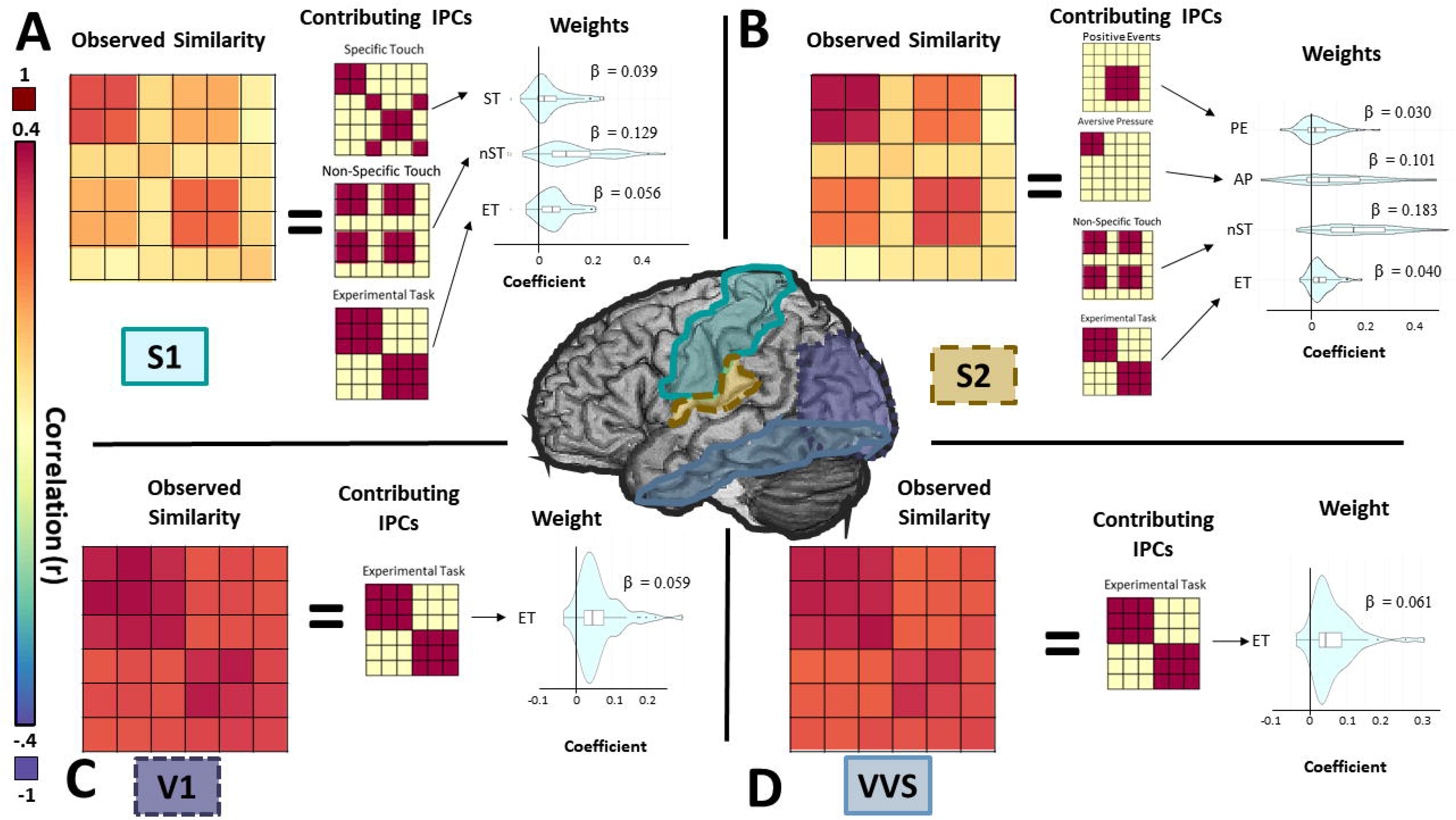
*Information pattern component in sensory cortices*. For illustrative purposes, this figure presents data from group sample analyses. A) Representational similarity in primary somatosensory cortex (S1) was characterized by IPCs indicating representations of discriminatory touch, including non-specific tactile salience (nST), and specific tactile experience (ST). B) Representation of both hedonic and discriminative tactile signals were observed in S2, with strongest representation of nST and aversive pressure (AP). C/D) Similarity of representations in primary visual cortex (V1) and ventral visual structures (VVS) were characterized by intra-task similarity, consistent with the conservation of visual stimuli within experimental tasks.

In consideration of the dominant contralateral input to S1 and the lateralized tactile stimulation across tasks (aversive pressure applied to the RIGHT thumbnail, and appetitive caress with a brush applied to the LEFT forearm), two additional PCM analyses were conducted in unilateral S1 ROIs. Importantly, these analyses added two additional theoretical pattern components, modeling the left and right lateralized components of non-hedonic tactile experience. For full details on IPC adjustments for unilateral PCM analyses, see Supplemental Material S2. In a pattern similar to that observed across the bilateral ROI, data from the random sample was predicted most strongly by components modeling non-specific aspects of tactile experience (Left S1: β_nST_ = 0.140; Right S1: β_nST_ = 0.129), with secondary contributions from components modelling experimental tasks (Left S1: β_ET_ = 0.052; Right S1: β_ET_ = 0.066). This similarity, however, was not observed for representations of non-specific tactile experience observed in the bilateral ROI. Left S1 did not represented aversive pressure as isolated from other forms of non-hedonic tactile states (β_AP_ = 0.032) and was lacking general representation for right lateralized non-hedonic touch. By contrast, right S1 represented non-hedonic touch experience (i.e., appetitive caress and scanner-generic touch as distinct states β_lST_ = 0.045). In both unilateral ROIs, components identified in the random sample significantly predicted the representational patterns of the held-out participants (Left S1: R^2^ = .237, *F* _(1,145)_ = 49.68, *p* < .001; Right S1: R^2^ = .240, *F* _(1,145)_ = 49.97, *p* < .001)

##### Secondary Somatosensory Cortex (S2)

For S2, the strongest predictor of representational patterns in the random samples was non-specific touch components (β_nST_ = 0.185). Additional components identified at a rate significantly greater than chance were aversive touch (β_AP_ = 0.094), experimental task (β_ET_ = 0.041), and the task-specific positive experience (i.e., caress, or safety; β_PE_ = 0.002). Combined, weighted components identified in the random sample accounted for an average of 41.1 % of the variance in the held-out participants (R^2^ = .411, *F* _(1,145)_ = 113.07, *p* < .001). This suggests that S2 may receive hedonic signals that are not represented in S1 (Figure 3B).

##### Visual Cortices

The most predictive pattern components in for both V1 and VVS representational patterns were found for components modeling task-related changes in experience, with no other pattern components identified at a rate significantly greater than chance. In V1, this component explained and average of 5.9% of the variance in the initial random sample, while in VVS, it explained an average of 6.1% of the variance. This weighted components, however, failed to significantly predict patterns observed in the held-out participants in cross-validation procedures (V1: R^2^ = .021, *F* _(1,145)_ = 4.27, *p* = .10; VVS: R^2^ = .028, *F* _(1,145)_ = 5.27, *p* = .07) This demonstrates that representational patterns in visual cortices may reflect visual attentional demands of the experimental task and are relatively uninformative of non-visual information, regardless of its hedonic value (Figure 3C/D). The inability to replicate significant findings in the held-out participants, however, indicates that activational patterns in these regions are likely not driven by information modeled in the current set of independent pattern components, but may instead be more accurately represented by pattern components modeling specific aspects of visual (rather than tactile) experience

#### Interoceptive regions of interest

##### Amygdalae

A combination of three pattern components were identified at a rate significantly greater than chance in the initial random sample for bilateral amygdalae. These components modeled a linear spectrum of tactile valence (β_TV_ = 0.014), non-specific tactile experiences (β_nST_ = 0.019), and global differences in experimental tasks (β_ET_ = 0.011). Pattern components identified in the random sample accounted for 11.6 % of the observed variance in the held-out participants (R^2^ = .116, *F* _(1,145)_ = 20.86, *p* = .0049). Notably, this regions housed representations of valence on a linear spectrum, where appetitive and aversive touch were most dissimilar – polar opposites of a shared representational space (Figure 4A).

**Figure 4.**
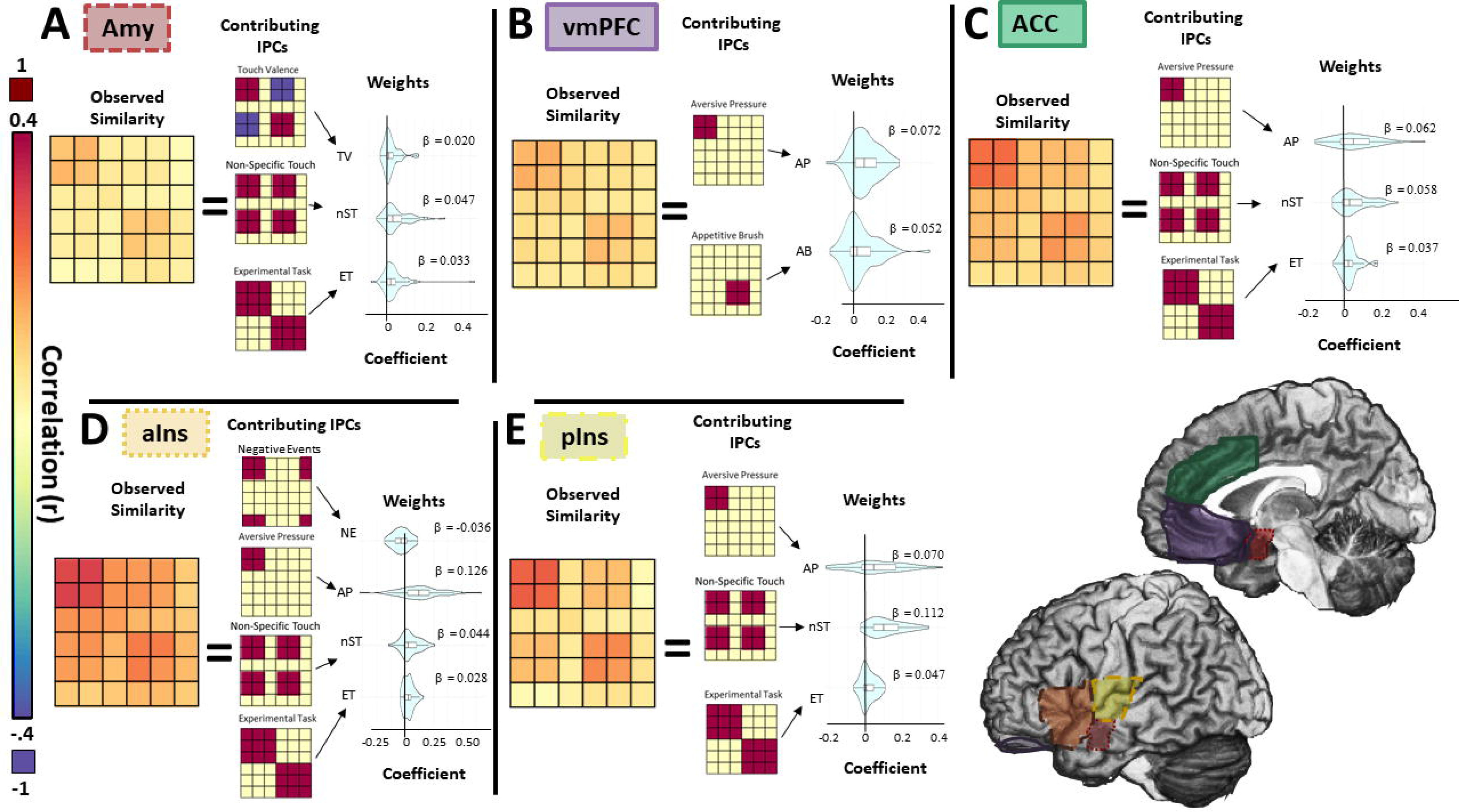
*Information pattern components in frontotemporal cortices*. For illustrative purposes, this figure presents data from group sample analyses. A) Amygdalae displayed patterns of activation consistent with representations of a unidimensional hedonic-tactile spectrum (TV), as well as tactile saliency and general task effects. B) Activation patterns in vmPFC were unique, with representation of appetitive and aversive tactile experience as dissociable contributing components. C/D/E) Anterior and posterior insula and ACC all displayed patterns of activation consistent with representation of tactile salience and aversive touch, though the specific biases for these types of information varied between the structures.

##### Ventromedial Prefrontal Cortex

The vmPFC representational patterns in the random samples were predicted most by a number of separate theoretical pattern components, most notably the two distinct hedonic touch components; ‘Aversive pressure (β_AP_ = 0.050)’ and ‘Appetitive caress (β_AB_ = 0.033)’. Additional components identified at a rate greater than chance in the random sample include those modeling differences in experimental task (β_ET_ = 0.011), non-specific tactile representations (β_nST_ = 0.015), and the temporal adjacency of experiences (β_TA_ = 0.005). Total variance accounted for in the held-out participant by the models identified in the random samples was on average 7.4 % (R^2^ = .074, *F* _(1,145)_ = 12.98, *p* = .023). This demonstrates that vmPFC activity contains information about the hedonic value of the tactile stimulation, representing positive and negative values as distinctly independent and non-opposing, signals (Figure 4B). Furthermore, the heterogeneity of non-hedonic component identification in this region this suggests that while vmPFC does consistently represents aversive pressure and appetitive caress representations, there may be extensive inter-participant variability for other processing in this region.

##### Anterior Cingulate Cortex

Aversive pressure (AP) was identified as the component with as the strongest predictor in the ACC data of the random samples (β_AP_ = 0.067). Additional components identified at a rate significantly greater than chance included those modeling non-specific tactile experience (β_nST_ = 0.055) and experimental task (β_ET_ = 0.034). Recombined, these components identified in the random samples predicted an average of 19.1 % of the variance in the held-out participants (R^2^ = .191, *F* _(1,145)_ = 37.84, *p* < .001). This suggests that ACC represents general tactile information but is particularly sensitive to tactile information associated with pain (Figure 4C).

##### Insula

The insula was anatomically subdivided at the anterior commissure into distinct non-overlapping anterior/posterior regions. Representational patterns in the anterior insula (aIns) were significantly predicted by four pattern components. In order of representational strength, these components modeled aversive tactile experience (β_AP_ = 0.107), non-specific tactile experience (β_nST_ = 0.050), experimental tasks (β_ET_ = 0.029), and task-specific negative-events (β_NE_ = −0.016). Components identified in the random sample predicted an average of 15.3 % of the variance in the held-out participants (R^2^ = .153, *F* _(1,145)_ = 28.74, *p* < .0023).

Similar to representations observed in aIns, in pIns, activity was significantly predicted by components modeling aversive tactile experience (β_AP_ = 0.059), non-specific tactile experience (β_nST_ = 0.102), experimental tasks (β_ET_ = 0.043). An additional representational component modeling tactile valence (β_TV_ = 0.009) was also identified in this region. In pIns, combinations of component identified in the random sample predicted an average of 36.3% of the variance in the held-out participants (R^2^ = .363, *F* _(1,145)_ = 90.82, *p* < .001). This demonstrates that whereas the general type of information processed across the insula may be similar for the anterior and posterior sections - each region sensitive to both hedonic and non-hedonic signals - the precise nature and dominance of these representations differ (Figure 4D/E).

### N-path Analyses

To assess the robustness of IPC contributions, a one-way ANOVA was conducted on the average number search paths required to find the best fitting component combination for each iteration (i.e., n-path data) which identified a significant main effect of region (F _(8,7992)_ = 195.507, *p* < .001). A follow-up series of independent sample t-tests (all reported p-values are Bonferroni-corrected) identified four distinct clusters of ROIs characterized by their n-path. A lower search path likely indicates either a poor fit of the IPC models (if only a single model is frequently identified; e.g., V1/VVS), or robust representations for a specific subset of models (if identified IPC > 1; e.g. vmPFC). By contrast, a higher n-path likely indicates more overlapping representational space (e.g., insular subdivisions). Visual areas required expansion of fewer paths than required by any other area (all *p* < .001), yet they did not differ significantly from each other (*p* = 1.0). vmPFC had more branching than either V1 or VVS but less than all other ROIs (all *p* < 0.001). ACC, amygdala, and S2 did not significantly differ from each other, yet required less expansion of search paths than S1 or either insular ROI. (all *p*s < .001). Finally, while the anterior insula did not significantly differ from either the posterior insula or S1 (both *p*s = 1), the posterior insula displayed greater branching than S1 (*p* < .001). The greater n-paths in these regions suggests that there is likely greater overlap of representational space in these regions between the modeled components compares to representations in other regions.

## Discussion

In this study we applied a novel form of pattern component modeling with representational similarity analysis to dissociate how discriminatory vs. hedonic tactile information, coded at the somatosensory receptors and carried by A- and C-/CT-fibers respectively, contribute to population code representations in the human brain. Distinct representations of hedonic information were observed in frontal and temporal structures, including ventromedial prefrontal cortex (vmPFC), insula (Ins) and anterior cingulate cortex (ACC), as well as in secondary somatosensory cortex (S2). Importantly, primary somatosensory cortex (S1) did not represent all tactile information coded by peripheral receptors. We did not observe any representation of positive hedonic touch signals carried by CT-fiber afferents, and only limited representation of negative hedonic signals carried by C-fiber afferents. Visual areas, including primary/secondary visual cortex (V1) and ventral visual structures (VVS), displayed no representation of either affective or discriminative touch information. By contrast, negative hedonic information made a minor contribution to representational patterns in S1 contralateral to the tactile stimulation, indicating that some nociceptive information may reach this area independent of frontotemporal processing.

Together, the findings support the hypothesis that sensations carried by hedonic-labeled tactile signals from C and CT-fiber pathways, despite their salience and homeostatic significance, are for the most part *not* represented in S1. Rather, this information is represented predominantly in frontotemporal structures more typically implicated in interoception (Craig, 2011; Pollatos et al., 2016; Strigo & Craig, 2016) and the central mediation of emotional relevance (McFarland & Sibly, 1975; Rolls, 2000; Todd et al., 2020). These findings suggest that that peripheral signals of positive and negative tactile experience are represented in frontotemporal structures, independent of non-hedonic touch; yet they lack distinct representation in early sensory cortices. Thus, tactile hedonic information is distinct from traditional exteroceptive signals highlighting non-traditional mechanisms (Kryklywy et al., 2020) by which prioritized information may be incorporated into emotionally-guided cognitive processes.

### Cortical representations for non-hedonic touch

In S1, neural activity displayed representational patterns that discriminated tactile experiences, as well as *non-valence specific* components, of tactile manipulations. The identification of representations of specific touch experiences (IPC: ST) in this area substantiates its traditional primary exteroceptive role in processing discriminatory tactile information as carried by out A-fiber afferents (McGlone & Reilly, 2010; McGlone et al., 2014). Note that ‘specific touch’ is defined such that representational patterns for trials with no tactile manipulation (i.e., the somatosensory experience generated by lying in a scanner) have equivalent strength to the representations of trials with tactile manipulation. Thus, its manifestation is unlikely to be generated by peripheral hedonic signalling alone, as non-hedonic tactile experience elicits its own representational state. S1 strongly represented the experience of ‘non-specific touch’ (IPC: nST), indicating activity in this region was driven by salient tactile experiences with a shared representational space for both appetitive and aversive tactile manipulations. This suggests that these representations are not shaped by information carried by C- and CT-fiber independently, as the two distinct peripheral signals are represented by overlapping activation patterns. These are potentially mediated by re-entrant projections of tactile salience from other frontotemporal structures (Pessoa & Adolphs, 2010; Vuilleumier, 2005). Non-specific touch representation is likely to be either a discriminatory representation of body location (i.e., arm; not dependent on C- or CT- fiber activation) or general tactile salience (may or may not integrate information from C- and CT-fiber activation; i.e., hedonic salience). In support of the latter interpretation, there is evidence that S1 likely integrates re-entrant hedonic signals from multisensory emotion-related regions (Orenius et al., 2017).

Components indexing unprocessed projections of hedonic-labeled afferent pathways (i.e., ‘aversive pressure’ and ‘appetitive caress’) were absent in bilateral representational patterns observed in S1. This suggests that these hedonic signals may not be instantiated in traditional somatosensory processing structures despite originating as external cutaneous sensation. Interestingly, unilateral investigation of left S1 did identify a representation pattern associated with aversive pressure, though this was observed in the absence of specific tactile discriminability (i.e., ST or rST). This indicates that discriminative and nociceptive information may be integrated prior to reaching S1 (for candidate regions, see Abraira et al., 2017; Marshall & McGlone, 2020; Neubarth et al., 2020). Alternatively, it may be that sustained changes in tonic firing of rates of slow adapting mechanoreceptors (for review, see Abraham & Mathew, 2019; Knibestol, 1975) in response to the strong pressure manipulation (right hand), result in a distinct tactile representation in left S1 during CS- trials (pressure task). This representation would be distinct from the representation of scanner-generic sensation (right hand) experienced during the caress (which was applied to the left hand). Given substantive evidence to support both interpretations, it is likely that the observed pattern component contributions reflect a combination of these processes; however, we found no evidence of unique representational patterns in S1 for signals of appetitive hedonic information carried by CT-fiber pathways.

Taken together, non-hedonic tactile representation, likely of signals carried along A-fiber pathways (McGlone & Reilly, 2010), appear to dominate activity in early somatosensory cortices. While some evidence for hedonic representations in these regions exists, it appears that pre-cortical integration of A and C-/CT-fibre pathways (Abraira et al., 2017; Marshall & McGlone, 2020; Neubarth et al., 2020), or re-entrant feedback from higher order integrative structures (Pessoa & Adolphs, 2010; Vuilleumier, 2005) are the most probable source of these representations.

### Cortical representations for hedonic touch

Amongst all regions investigated, only vmPFC displayed independent representation of both appetitive and aversive touch (IPCs: AC/AP). This suggests that this region either A) receives information carried along C- and C-tactile fiber afferents as distinct signals prior to their integration with each other or other tactile information, or B) has decomposed an integrated hedonic representation back into distinct signals of positive and negative value to inform situation specific behaviours and decision. The potential of first order representation of peripherally labeled hedonic signals in vmPFC is particularly intriguing considering the critical role these ventral structures play in appraising emotional salience to guide value-based decision making (Dixon et al., 2017; Euston et al., 2012; Hiser & Koenigs, 2018). Propagation of hedonic-labeled tactile signals to these regions independent of any prior cortical processing would act as a mechanism to facilitate the prioritization of evolutionarily relevant sensation (Kryklywy et al., 2020), and allow for expedited integration of action-outcomes into value appraisal to guide decision-making processes.

The absence of distinct representations for pleasurable tactile signals in both the anterior and posterior insula is notable, as these regions have been highlighted as potential cortical recipients of C-tactile fiber signalling (Olausson et al., 2002; Rolls et al., 2003). While this may be due to variability in response to the appetitive touch manipulation used in the current design, the identification of a clear appetitive caress pattern component in the vmPFC indicated that this is unlikely. Instead, it may be that aversive touch is the more salient tactile signal for the immediate well-being of an organism (Rolls, 2000), and thus given greater priority of resources. Furthermore, much of the prior work identifying modulation of insula activity by pleasurable touch has been performed either independent of aversive touch (Olausson et al., 2002), or treating the two signals orthogonally without direct comparison (Rolls et al., 2003). This leaves open the possibility that previous results were driven by general affective salience of the tactile cue (as observed in the current work), rather than pleasurable sensation alone. Related to this, additional consideration should be given to the potential for sensory adaptation or habituation to hedonic tactile experience (McBurney & Balaban, 2009; Morrison, 2016). The current design relies on the averaged representational similarity across six distinct tactile exposures for each condition. Thus, it is possible that differences in the time course of downregulating tactile signalling between painful and pleasurable stimulation following multiple exposures may underlie some of the differences in representational strength of hedonic information. This question could be targeted by future studies that dissociate early and late exposure to hedonic touch.

Though distinct representation of appetitive touch was identified only in the vmPFC, distinct representations of aversive pressure were identified within the ACC as well as the anterior and posterior insula. Notably, both of these regions are heavily implicated in the representation of painful experience (Corradi-Dell’Acqua et al., 2016; Kragel et al., 2018) and are postulated to underlie awareness of one’s own internal homeostatic balance (Craig, 2011, 2015; Pollatos et al., 2016; Strigo & Craig, 2016), characterized as the *interoceptive self* (Craig, 2015). One potential explanation for this pattern of results is that hedonic-labeled peripheral afferents are not processed as tactile signals in the traditional view of sensation (Gazzaniga et al., 2019; Pinel & Barnes, 2018). That is, they may not be instantiated in neocortex as representing the experience of contact with external objects in the environment. Rather, information carried along these pathways indicates internal concerns about homeostatic threat or social safety (Craig, 2011, 2015) and manifest cognitively as emotional feelings congruent with these states. Such patterns of representation demonstrate a potential dual function of proximal senses, consistent with Sherrington’s classic distinction between exteroception and interoception: they represent not signals of the external world, but instead of the body itself, Sherrington’s material “me” (Sherrington, 1906). Information about the internal state acts can then act as an immediate mechanism for motivating response, independent of its representation as an exteroceptive tactile experience or any other form of cognitive processing.

### Integrated representation of tactile experience

Multiple structures displayed patterns of activity that indicated representation of C- and CT-fiber afferent information, though not in a mutually exclusive manner. Rather, representational patterns showed distinct similarity/dissimilarity between hedonic conditions, indicative of prior processing and representational integration of this information. Specifically, in the ACC as well as the anterior and posterior insula, patterns of neural activity were found to represent non-specific touch (IPC: nST) in addition to aversive pressure (IPC: AP). While the representation of non-specific tactile experiences is possibly dependent on information integrating both types of unmyelinated hedonic tactile afferents (C- and CT- fibers), the similarity of representation between the two valence manipulations indicate it is unlikely to be a first order representation of information carried along these fibers. Rather, this integrated representations likely indicate processing of sensory information prior to their affective representations, in a manner more typical of traditional models for emotional prioritization (Pessoa & Adolphs, 2010; Rolls, 2000, 2019; Vuilleumier, 2005).

While none of the anterior or posterior insula, ACC, or vmPFC were found to display opposing representations of hedonic valence, in the amygdala, a clear representation of the hedonic experience of touch valence (IPC: TV) was observed. Here, signals of tactile valence were represented as a single linear vector, with hedonic conditions represented as polar ends of a single valence spectrum: pleasure on one end and pain on the other. Thus, prior to its representation, or as part of its processing, in the amygdala, tactile information initially carried as unique signals along non-overlapping sensory afferents must be integrated into the same representational space. These amygdalar bi-polar valence representations are consistent with those identified in the olfactory domain (Jin et al., 2015), but have not been observed for either gustatory or visual hedonic information (Chikazoe et al., 2014, 2019). This divergence indicates a probable modal-specificity of hedonic processing in the amygdala rather than a centralized a-modal representation of emotional information (Miskovic & Anderson, 2018). In the current work in particular, this unidimensional hedonic vector may be related to the association of the tactile sensation with the concurrent visual stimuli rather than with the raw tactile signals in isolation, as multiple studies in both humans and non-human primates have implicated this region in guiding affect-biased attention (Todd et al., 2020) and emotional learning in vision (Everitt et al., 2003; Morris et al., 1998). Of note, however, there is substantial evidence to suggest that both hedonic responding and attentional biases may be heavily influenced by individual differences between subjects (Harjunen et al., 2017; Nielsen et al., 2009; Todd et al., 2012), in addition to researcher-expected hedonic experience. Future work conducted in a larger sample population could provide additional insight into this possibility.

## Conclusion

Somatosensation, from the point of initial neural encoding, contains more information about the environment than traditionally believed. Beyond discriminative information pertaining to the identity of objects contiguous to ourselves, this system also signals the value of an object to our wellbeing — the pain and pleasure of contact with it. In the current study, we performed a novel theory-driven implementation of multivariate pattern component modeling to deconstruct observed representational patterns into discrete contributing components. Using this approach, we demonstrated that hedonic tactile information is not processed in the same fashion as non-hedonic tactile information. The full spectrum of hedonic tactile information is not uniquely represented in primary somatosensory cortices but is represented in frontotemporal structures. Notably, representations of hedonic tactile information are observed in brain areas proposed to underlie the emergence of an interoceptive self and the capacity for cognitive decision making. We propose that somatosensory signaling contains two distinct potential channels for affective prioritization. One channel, propagating through somatosensory cortex to frontotemporal regions processes the integrated experience of tactile sensation, extracts valuable information about the associated hedonic values. The second, potentially bypassing early somatosensory structures in favour of traditionally integrative regions, does not extract hedonic information, but rather uses what is already present in peripheral channels to inform representation our own homeostatic representations and guide our decision-making processes. Through this lens, somatosensation, originating from cutaneous mechanoreceptors through contact with the external world, is not only critical for our exteroception – *feeling about the world around us*. but also for our interoception and emotion – *feelings about the world inside*.

## Methods

### Participants

Four-hundred and eighty-eight participants were recruited from Cornell University to complete an initial behavioural pilot assessment of affiliative responding to tactile stimulation. Of these, 107 participants (x□_age_ = 21.1, *sd* = 2.8; 41F) were recruited to complete the current study. We were unable to complete preprocessing of data for 40 participants: 27 participants had raw data corrupted related to server-transfer errors prior to preprocessing, for five we were unable to obtain convergence during the multi-echo independent component analysis (ICA), five did not have correct stimulus timing information, and three were excluded due to motion artifacts. Results from the remaining 67 participants are reported. All participants gave written, informed consent and had normal or corrected-to-normal vision. Participants were pre-screened for a history of anxiety and depression as well as other psychopathology, epilepsy and brain surgery. Pre-screening was followed up in person by an additional interview to ensure inclusion criteria were met. As this study was conducted as part of larger research program, all participants provided saliva samples for genotyping, and fecal sample for microbiome analyses. The experiment was performed in accordance with the Institutional Review Board for Human Participants at Cornell University.

### Stimuli and Apparatus

Three male and three female faces with neutral expressions were chosen from the Karolinska directed emotional faces picture set (Goeleven et al., 2008). These faces were used as conditioned stimuli (CS) in two classical conditioning paradigms, each containing two CS+ and one CS- stimuli. Unconditioned stimuli (US) consisted of either aversive pressure delivered to the right thumb, or appetitive caress to the participant’s left forearm. These tactile manipulations were aimed to maximally activate C- and C-tactile somatosensory afferent respectively. Aversive pressure stimuli were delivered using a custom designed hydraulic device (Giesecke et al., 2004; Lopez-Sola et al., 2010) capable of transmitting controlled pressure to 1 cm^2^ surface placed on the subjects’ right thumbnail. Applied pressure levels were individually calibrated for each participant prior to the experiment to ensure that the pressure intensity was experienced as aversive but not excessively painful. Light appetitive caress lasting ~4 s were manually applied to the left forearm with a brush by a trained experimenter to maximally activate CT-fiber pathways (McGlone et al., 2014). Only participants who had previously demonstrated reliable affective responses to the tactile manipulations in the initial behavioural pilot (e.g., positive response to caress; see below for details) were invited to participant in the scanning session, as this indicated likely recruitment of C- and CT- fibre pathways of the hedonic tactile manipulations (which can be subject to inter-participant variability).

### Procedure

While undergoing functional MR scanning, participants completed two separate conditioning tasks (appetitive conditioning and aversive conditioning), each involving a series of tactile and visual pairings (Figure 1A) (Visser et al., 2015). In each task, participants completed seven CS-only blocks interleaved with six CS-US paired blocks. Single blocks of either the CS-only or the CS-US pairing contained one presentation of each facial stimulus (i.e., 3 face stimuli, 2 CS+ and 1 CS-, per block of each conditioning task). Individual trials consisted of an initial fixation period (19500 ms) followed by the presentation of a face (4000 ms). A fixed and long interstimulus interval (19500 ms) was included in the experimental design to reduce intrinsic noise correlations and enable trial by trial analyses by means of RSA (Visser et al., 2016; Visser et al., 2013). During CS-only trials, all faces were presented without tactile stimulation. During CS-US paired trials, two of three facial stimuli presentations overlapped with tactile stimulation, thus creating two CS+ and one CS-. The US was delivered from the midpoint of the face presentation (2000 ms post-onset), remained for the rest of the time the face was visible (2000 ms) and persisted following the offset (2000 ms; total US = 4000 ms). The order of face presentation was randomized within each CS-US paired block. Participants completed two experimental tasks (one for each US, order counterbalanced across participants), totaling 26 blocks (6 CS-US paired and 7 CS only blocks for each US type).

Note that this paradigm was designed to target two distinct questions: 1) How and where are discriminatory and hedonic tactile signals represented in the brain? and 2) How do novel affective associations change the neural representation of conditioned stimuli? The current work focuses on the former question. Importantly, trials described as CS- during CS-US pairings are not independent of tactile stimulation. Rather, these trials lacked pain or brush stimulation, but participants still experienced tactile stimulation from the scanner, their clothes etc.

### MRI Acquisition and Preprocessing

MR scanning was conducted on a 3 Tesla GE Discovery MR scanner using a 32-channel head coil. For each subject, a T1-weighted MPRAGE sequence was used to obtain high-resolution anatomical images (TR = 7 ms, TE = 3.42 ms, field of view (FOV) 256 × 256 mm, slice thickness 1 mm, 176 slices). Functional tasks were acquired with the following multi-echo (ME) EPI sequence: TR = 2000 ms, TE1 = 11.7 ms, TE2 = 24.2 ms and TE3 = 37.1 ms, flip angle 77°; FOV 240 × 240 mm. These parameters are consistent with recent work demonstrating improved effect-size estimation and statistical power for multi-echo acquisition parameters (Lombardo et al., 2016). Specifically, the multi-echo sequence was chosen due to its enhanced capacity for differentiating BOLD and non-BOLD signal (Kundu et al., 2012; Kundu et al., 2014), as well as its sensitivity for discrimination of small nuclei in areas susceptible to high signal dropout (Markello et al., 2018). A total of 468 volumes, (102 slices, thickness 3.5mm; 72 × 72 acquisition matrix, 3.33 mm X 3.33 mm) were acquired for each functional run. Pulse and respiration data were acquired with scanner-integrated devices.

Preprocessing and analysis of the fMRI data was conducted using Analysis of Functional NeuroImages software (AFNI; Cox, 1996) and the associated toolbox *meica.py* (Kundu et al., 2014; Kundu et al., 2017). For maximal sensitivity during multivariate pattern detection, no spatial smoothing was performed on the data (Haynes, 2015). Preprocessing of multi-echo imaging data followed the procedural steps outlined by Kundu et al. (Kundu et al., 2013; Kundu et al., 2012), and used independent component analyses to define a set of components using their TE dependence to be individually classified as BOLD or non-BOLD (e.g., motion artifact). An optimally combined (OC) dataset was generated from the functional multi-echo data by taking a weighted summation of the three echoes, using an exponential T2* weighting approach (Posse et al., 1999). Multi-echo principal components analysis (PCA) was first applied to the OC dataset to reduce the data dimensionality. Spatial independent components analysis (ICA) was then applied and the independent component time-series were fit to the pre-processed time-series from each of the three echoes to generate ICA weights for each echo. These weights were subsequently fitted to the linear TE-dependence and TE-independence models to generate F-statistics and component-level κ and ρ values, which respectively indicate BOLD and non-BOLD weightings. The κ and ρ metrics were then used to identify non-BOLD-like components to be regressed out of the OC dataset as noise regressors. Regressor files of interest were generated for all individual trials across the experiment, modelling the time course of each stimulus presentation during each run (36 total events: 2 tasks X 6 CS-US blocks X 3 CS, with each event beginning at the face presentation onset). The relevant hemodynamic response function was fit to each regressor for linear regression modeling. This resulted in a β coefficient and t value for each voxel and regressor. To facilitate group analysis, each individual’s data were transformed into the standard brain space of the Montreal Neurological Institute (MNI).

### fMRI Analyses: Structural Regions of Interest

To assess tactile (pressure *and* caress *and* non-specific touch) and hedonic (pressure *vs.* caress) representations in neural patterns, nine bilateral regions of interest (ROIs) were generated from the standard anatomical atlas (MNIa_caez_ml_18) implemented with AFNI. Selected ROIS were: Primary somatosensory cortex (S1), secondary somatosensory cortex (S2), primary/secondary visual cortex (V1), ventral visual structures (VVS), amygdala, ventromedial prefrontal cortex (Posse et al., 1999), anterior cingulate cortex (ACC) and separate posterior/anterior insula (Ins) divisions (consistent with its functional and histological divisions; for review, see Nieuwenhuys, 2012). S1 and V1 were selected as the primary sites of tactile and visual information respectively. VVS were chosen due to their role in visual classification (Kanwisher et al., 1997; Kravitz et al., 2013). Amygdala, vmPFC, ACC and posterior/anterior Ins divisions were selected for their hypothesized roles in affect and pain representations subdivisions (Anderson & Phelps, 2002; for rationale behind multiple insular ROIs, see Cauda et al., 2012; Chikazoe et al., 2014; Kragel et al., 2018; Orenius et al., 2017) For extended details on defining our ROI see Table 1.

### fMRI Analyses: RSA and Ideal Model Specification

In order to identify and compare the representational pattern elicited by the experimental conditions, representational similarity analysis (RSA; Kriegeskorte & Kievit, 2013; Figure 1B; Mur et al., 2009) was performed using the PyMVPA Python package (Hanke et al., 2009). For each participant, a vector was created containing the spatial patterns derived from β coefficients from each voxel related to each particular event in each ROI. Pairwise Pearson coefficients were calculated between all vectors of a single ROI, thus resulting in a similarity matrix containing correlations for all trials for each participant (i.e., how closely the pattern of voxel activation elicited in one trial resembles the patterns of voxel activation observed in all other trials). Fisher transformations were performed on all similarity matrices to allow comparisons between participants. Correlation matrix transformations were performed using Matlab (The MathWorks, Natick, Massachusetts, USA) and BIC analyses were conducted in R (RCoreTeam, 2013) with the package PCMforR (J. H. Kryklywy et al., 2021).

A novel theory-guided implementation of pattern component modelling (PCM; Kriegeskorte and Kievit, 2013; Diedrichsen et al., 2018 was performed using thirteen predefined models of potential information pattern component (IPCs). These models were generated to represent similarity matrices that would be observed in the experimental data if it were to contain perfect representation of distinct sources of information. IPCs were constructed for 1) Experimental task, 2) Non-specific touch, 3) Specific touch, 4) Appetitive caress 5) Aversive pressure, 6) Touch valence, 7) Positive events, 8) Negative events, 9) All valence, 10) Salience, 11) Face Stimulus, 12) Violation of expectation, and 13) Temporal adjacency. For a description of all thirteen IPCs, see Table 2 and for an extended description see Supplemental Material S1.

To determine the IPC combinations that best explained the observed correlations in the data for each ROI, we conducted Monte-Carlo cross validated PCM using Bayesian Information Criterion (BIC) to fit our pattern component models. As many the theoretical independent pattern components (IPCs) in the current experiment contained some overlapping information, a regression of all potential components would be insufficient to identify those most informative. To address this, an uninformed *greedy best-first search* (*GBFS*) algorithm (Doran & Michie, 1966) was implemented to identify the best fitting IPC combination in a step wise manner (see Supplemental Figure SF1). This allows us to distinguish between subtle nuances in representational patterns by iteratively building up our identified represented components in a step-wise manner rather than all at once, thus comparing the fit of overtly similar component patterns both independent from, and in combination with, each other. Initial model testing was conducted by fitting of each independent IPC to the observed similarity for a given ROI (Level 1). Upon identification of the best fitting IPC (IPC_B1_), model fitting was conducted on each remaining IPC in combination with IPC_B_ (Level 2). The IPC combination (i.e., IPC_B1_ + IPC_B2_) that provided the best fit to the ROI data would be held as a constant for model fitting in Level 3. This process was repeated iteratively until no addition of remaining IPCs led to an improved fit to the ROI. A ΔBIC > 2 was defined as indicative of an improved fit (Fabozzi, 2014). Following similar equivalency criteria, all IPC combinations at a given search level with ΔBIC scores < 2 to the best fitting combination were also extended to path completion (Fabozzi, 2014). This approach allowed for the decomposition of observed representational patterns into multiple unique contributing sources of information. The independent pattern components identified as contributing to the representational space were subsequently fit using linear regression modeling to the observed similarity in the original ROI (see Supplemental Figure SF1).

To ensure that regression fits were not a product of overfitting, these analyses were performed as a cross validation procedure. Initial model fitting was performed on a randomly selected sample of participants (‘*random-sample’*, RS = 60), with the identified components fit as a predictor to data from the remaining participants held-out of this initial sample (the ‘*hold-out’:* HO = 7). Monte-Carlo cross-validation (CV; Picard & Cook, 1984) parameters were chosen to maximize CV performance by minimizing CV-variance while maximizing model selection accuracy (Arlot & Celisse, 2010). These analyses identified IPCs contributing to representational patterns for each ROI in the RS. Beta coefficients and intercepts, determined by fitting these IPCs as predictors to the experimental data, were used to create a reconstructed and averaged dataset. The reconstructed dataset was then fitted as a predictor to the HO, with each iteration approximating a single fold of a 10-fold validation.

Specific outputs of interest included the proportion of CV iterations (i = 1000) in which an IPC was identified as a contributing component of the experimental data in the RS, the average weight of representation for significantly identified components to the RS, the number of search paths required to fir the data on each iteration (n-path) and the model fit of the reconstructed RS data to the HO (Figure 2). Proportion of iterations for IPC identification were compared to chance identification for each ROI (i.e., # of IPCs identified / total # of IPCs).

