## Supplemental Material for "Decoding representations of discriminatory and hedonic information during appetitive and aversive touch"

**Supplementary Material**

### *S1. Information Pattern Component (IPC) Extended Descriptions*

Pattern component modelling (PCM), an approach foreshadowed by Kriegeskorte (2013) and outlined by Diedrichsen et al (2018), was adapted and implemented in the current study. *Information pattern components* (IPCs) were created to describe the similarity matrices that would be observed if the data were to ideally represent a single type of information perfectly. It should be noted that each 'correlation' in these matrices is the average value of all correlations between relevant trial types. The average correlation within each condition (i.e., appetitive CS+1 with appetitive CS+1) is the mean of 30 individual correlations (matrix of 6 exposures with auto-correlations removed), while the average correlation between conditions (e.g., appetitive CS+1 with appetitive CS-) is the mean of 36 individual correlations (matrix of 6 exposure to each condition).

Note that the IPCs were designed to capture information about the specific manipulations of the current experiment, with a particular focus on potential representations of tactile experience. Thus, they may not accurately reflect information represented in brain regions processing predominantly non-tactile information.

#### *Experimental task (ET)*

'Experimental task' was conceptualized as a representational overlap for all trials contained in one conditioning task (i.e., the aversive vs appetitive condition task). This included both CS-US paired and CS-only (CS-) stimuli for the conditioning task. Ideal representation of 'Experimental task' was defined as perfect correlation ($r = 1$) between all trials in each specific conditioning task. All correlations for trial across conditioning tasks were set to $r = 0$.

#### *Non-specific touch (nST)*

'Non-specific touch' was conceptualized as identical representations for all trials where a tactile manipulation occurred. Importantly, this IPC does not represent any discriminable information between the two tactile manipulations, but rather defines them as sharing representational space. This IPC may be derived from information carried by non-hedonic (A fiber) or hedonic (C/C-Tactile) peripheral channels; non-hedonic signals may indicate quality/strength of tactile information while hedonic signals may indicate emotional saliency of tactile experience. Note, however, that this pattern is unlikely to occur as a result of first order hedonic projection (which are separated into unique positive and negative valence signals), as it includes, by definition, a shared representational space across conditions.

Ideal representation of nST was modeled as perfect representational overlap (r = 1) between appetitive brush-paired trials and other appetitive brush (AB-AB) trials, aversive pressure-paired trials and other aversive-pressure (AP-AP) trials, as well as aversive pressure and appetitive brush (AP-AB) trials. All correlations involving no tactile manipulations were defined as r = 0.

*Specific touch (ST)*

‘Specific touch’ was conceptualized as representing each tactile experience as occupying a unique representation space. Critically, in addition to a unique representational space for appetitive and aversive experience, this IPC also defines the tactile experience of no active manipulation as occupying its own representation space. Thus, a pattern similar to this IPC cannot be derived from hedonic tactile channels, but rather must rely on alternate signals of discriminative touch (i.e., information carried by A fibers). Ideal representation of ‘Specific touch’ entailed perfect correlation (r = 1) between all trials in which similar tactile experience occurred. This included AP-AP trials, AB-AB trials, and no manipulation (i.e., generic scanner sensation) trials. All additional correlations crossing tactile experience types were set to r = 0.

*Appetitive brush (AB)*

‘Appetitive brush’ was conceptualized as representing information carried by C-tactile fiber pathways and the associated experience of pleasant touch. Ideal representation of ‘Appetitive brush’ was defined as perfect correlation (r = 1) between all trials that involved the delivery of an appetitive brush stroke to the participant’s arm (i.e., AB-AB trials). It should be noted that similarity between these trials could be ascribed to either the tactile sensation of brushing or the hedonic nature of the experience. All additional correlations were set to r = 0.

*Aversive pressure (AP)*

‘Aversive pressure’ was conceptualized as representing information carried by C- fiber nociceptive pathways and the associated experience of painful stimulation. Ideal representation of ‘Aversive pressure’ was defined as perfect correlation (r = 1) between all trials that involved the delivery of aversive pressure to the thumbnail (i.e., AP-AP trials). As with Appetitive brush, it should be noted that similarity between these trials could ascribed to either the tactile sensation of pressure or the hedonic nature of the experience. All additional correlations were set to r = 0.

*Touch valence (TV)*

‘Touch valence’ represents painful and pleasant tactile experience as opposite ends of a shared linear representational space. Ideal representation of ‘Touch valence’ was defined as a linear representation of the hedonic aspect of the tactile manipulation. For this, all correlations between all AB-AB trials and all AP-AP trials were set to r = 1. To represent the contrasting hedonic experience of AB-AP trials, these correlations were set to r = -1. All additional correlations were set to r = 0.

*Positive events (PE)*

‘Positive events’ were conceptualized as a general representation for individual trials that are positively valenced relative to the context in which they are situated. This includes both the positive valence related to the *presence* of pleasant brushing during the appetitive conditioning task and the *absence* of painful pressure during the aversive conditioning task. Ideal representation of ‘Positive events’ was defined as perfect correlation (r = 1) between all trials where the event was positive relative to its context. Positive trials (PT) included the delivery of a brush stroke during the appetitive conditioning task as well as the absence of pressure in the aversive conditioning task. All additional correlations were set to r = 0.

*Negative events (NE)*

‘Negative events’ were conceptualized as a general representation for individual trials that negatively valenced relative to the context in which they are situated. This includes both the feeling of painful pressure during the aversive conditioning task, as well as the absence of pleasurable brushing during the appetitive conditioning task. Ideal representation of ‘Negative events’ was defined as perfect correlation (r = 1) between all trials where the event was negative relative to the rest of the conditioning task. Negative trials (NT) included the delivery of pressure during the aversive conditioning task as well as the absence of a brush stroke in the appetitive conditioning task. All additional correlations were set to r = 0.

*All valence (AV)*

'All valence' was conceptualized as a linear representation of the hedonic aspects of the experimental procedure relative to the task in which they were situated. Here 'Positive events' and 'Negative events' are independent IPCs that are considered to be opposite ends of a shared linear representational space. To represent this, all PT-PT and NT-NT correlations were set to r = 1. All PT-NT correlations were set to r = -1.

*Salience (Sa)*

'Salience' was conceptualized as a linear representation of the salience of individual trials, with 'high salience' and 'low salience' considered as opposite ends of a shared linear representational space. For this, correlations between all highly salient trials (i.e., AB and AP trials) were set to r = 1. In addition, correlations between all minimally salient trials (i.e., CS- trials) were also set to r = 1. All correlations between highly and minimally salient trials were set to r = -1.

*Face stimulus (FS)*

'Face stimulus' was conceptualized as a unique representational space for each independent facial identity. To model this, FS defines correlations for all trials contained an identical visual CS (6 in total, 3/experimental task) as r = 1. All additional correlations between different CS were set to r = 0.

*Violation of expectation (VE)*

Violation of expectation was conceptualized as a shared representation space for the less likely tactile outcome in each conditioning task As there were two CS-US paired trials for each CS-only trials for each experimental task, the less probable outcome was always the CS-only trials. As such, the VE IPC modeled all CS-minus-CS-minus correlations (within and across experimental task) as r = 1. All additional correlations were set to r = 0.

*Temporal adjacency (TA)*

‘Temporal adjacency’ was conceptualized as representing any task-irrelevant cognitive processes that may extend beyond single modeled events, and thus inform the representational pattern of temporally adjacent trials. Ideal representation of ‘Temporal adjacency’ was defined as perfect correlation ($r = 1$) between all comparisons that included trials that were temporally contained within the same block (i.e., within each group of three CS presentations). Due to the removal of autocorrelation from within-condition averaging, these did not contain temporally adjacent trials, nor did any correlation of trial between experimental tasks. Thus, correlations for both of these comparisons were set to $r = 0$.

#### ***S2. Unilateral IPC descriptions for additional S1 analyses***

*Right specific touch (rST)*

This IPC represents the discriminable tactile experience expected elicited from the right side of the body. It represents both aversive pressure pain (applied to the right thumb) and generic scanner touch (including that occurring on the right side during appetitive brush trials) as dissociable tactile experiences. This IPC is predicted to manifest in representation in left S1.

*Left specific touch (lST)*

This IPC represents the discriminable tactile experience expected elicited from the left side of the body. It represents both appetitive brushing (applied to the left forearm) and generic scanner touch (including that occurring on the left side during the application of aversive pressure) as dissociable tactile experiences. This IPC is predicted to manifest in representation in right S1.
