## Supplemental Figures for "Decoding representations of discriminatory and hedonic information during appetitive and aversive touch"

### Supplementary Figure SF1

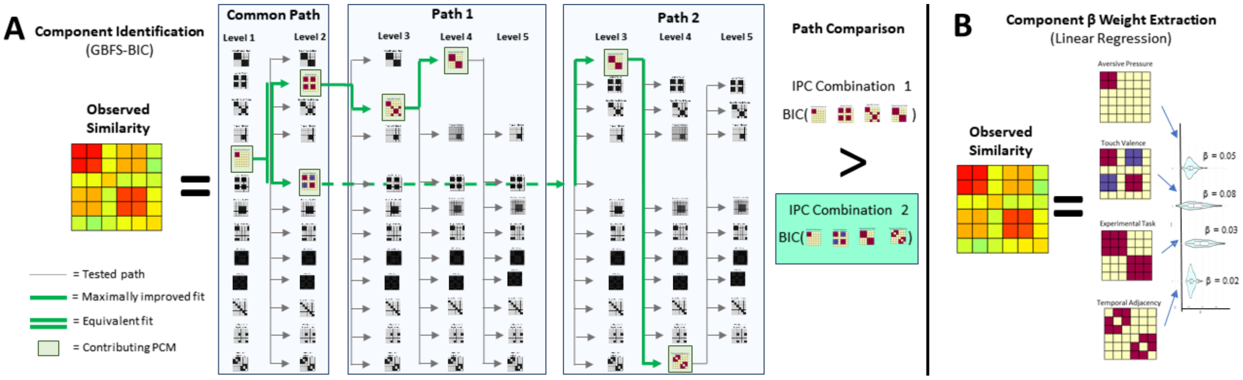


**SF1:** *PCM and component weighting. A) Information Pattern Component (IPC) Identification.* A greedy best-first search algorithm (GBFS) with Bayesian Information Criterion (BIC) was used to identify the combination of IPCs that best predicted experimental data. Level-1 search independently fit thirteen unique IPCs to observed representation similarity. The IPC with optimal fit was combined with all remaining models (Level-2) to determine the combination of IPC represented in the ROI. This process was done iteratively until the addition of none of the IPCs improved model fit. B*) IPC Weighting.* Multiple regression allowed extraction of β value indicating the representational strength for each contributing IPC.
