## Supplemental Tables for "Decoding representations of discriminatory and hedonic information during appetitive and aversive touch"

**Supplementary Table ST1:** *Regions of Interest*

| Region | Abbr. | Traditional Processing | Volume ($mm^3$) | Atlas* | Label** |
|---|---|---|---|---|---|
| Primary somatosensory cortex | *S1* | Sensory | 60 200 | MNIa_caez_ml_18: | postcentral gyrus |
| Left S1 | *lS1* | Sensory | | MNIa_caez_ml_18: | left postcentral gyrus |
| Right S1 | *rS1* | Sensory | | MNIa_caez_ml_18: | right postcentral gyrus |
| Secondary somatosensory cortex | *S2* | Sensory | 12 981 | MNIa_caez_ml_18: | L/R rolandic operculum (posterior to the anterior commissure; y > 0) |
| Primary/Secondary visual cortex | *V1* | Sensory | 76 968 | MNIa_caez_ml_18: | lingual gyrus + calcarine gyrus + cuneus (y > 60) |
| Ventral visual structures | *VVS* | Sensory | 105 664 | MNIa_caez_ml_18: | inferior temporal gyrus + inferior occipital gyrus + fusiform gyrus |
| Amygdalae | *Amy* | Affect | 3 664 | MNIa_caez_ml_18: | amygdala |
| Ventromedial prefrontal cortex | *vmPFC* | Affect Cognitive | 42 160 | MNI_vmPFC: | entire atlas |
| Anterior cingulate cortex | *ACC* | Sensory Affect Cognitive | 26 528 | MNIa_caez_ml_18[a:] MNI_vmPFC[b:] | [anterior cingulate + middle cingulate][a] minus overlap[b] (anterior to the anterior commissure; y < 0) |
| Insula (anterior) | *aIns* | Affect Interoceptive | 17 728 | MNIa_caez_ml_18: | insula lobe (anterior to the anterior commissure; y < 0) |
| Insula (posterior) | *pIns* | Sensory Affect | 8 296 | MNIa_caez_ml_18: | insula lobe (posterior to the anterior commissure; y > 0) |

***** All atlases were transformed to MNI space prior to their implementation or manipulation

** All labels refer to the bilateral structures, unless otherwise indicated

**Supplementary Table ST2:** *GBFS*-BIC Analysis Paths

| | | | | **Information Pattern Component** – BIC Score | | | | | | | | | | | | |
|---|---|---|---|---|---|---|---|---|---|---|---|---|---|---|---|---|
| **ROI** | **None** | **All** | **Included+** | **ET** | **nST** | **ST** | **AC** | **AP** | **TV** | **PE** | **NE** | **AV** | **Sa** | **FS** | **VE** | **TA** |
| **S1** | -1034.08 | -1389.76 | - | -1119.00 | -1319.74 | -1195.35 | -1120.78 | -1188.25 | -1142.20 | -1036.30 | -1038.53 | -1062.32 | -1231.40 | -1128.35 | -952.898 | -1027.03 |
| | | | nST+ | -1405.82 | - | -1386.70 | -1322.49 | -1356.94 | -1398.61 | -1320.73 | -1323.06 | -1344.03 | -1319.99 | -1359.95 | -1198.08 | -1327.22 |
| | | | nST+ET+ | - | - | -1413.71 | -1398.70 | -1410.78 | -1412.63 | -1401.79 | -1403.37 | -1412.35 | -1401.70 | -1402.46 | -1269.41 | -1402.46 |
| | | | nST+ET+ST+ | - | - | - | -1408.15 | -1413.60 | -1410.12 | -1406 | -1407.64 | -1409.97 | -1410.12 | -1406.57 | -1275.21 | -1406.57 |
| **S2** | -779.51 | -1455.63 | - | -838.18 | -1375.29 | -938.8 | -867.218 | -1054.75 | -876.23 | -776.04 | -784.64 | -797.25 | -1100.80 | -862.85 | -781.56 | -772.60 |
| | | | nST+ | -1441.22 | - | -1415.80 | -1368.06 | -1450.19 | -1441.93 | -1370.53 | -1380.51 | -1389.95 | -1369.10 | -1394.69 | -1253.89 | -1386.84 |
| | | | nST+AP+ | -1476.14 | - | -1457.40 | -1463.51 | - | -1463.51 | -1464.24 | -1458.57 | -1445.33 | -1444.07 | -1451.23 | -1312.29 | -1453.43 |
| | | | nST+AP+ET+ | - | - | -1470.08 | -1470.43 | - | -1470.43 | -1479.58 | -1477.12 | -1469.88 | -1469.05 | -1468.89 | -1336.99 | -1468.89 |
| | | | nST+AP+ET+PE | - | - | -1472.44 | -1473.87 | - | -1473.87 | - | -1476.96 | -1476.96 | -1472.41 | -1473.82 | -1340.98 | -1473.82 |
| **V1** | -579.77 | -564.89 | - | -604.03 | -576.78 | -590.49 | -576.50 | -588.58 | -594.09 | -573.30 | -573.92 | -576.08 | -576.98 | -582.91 | -502.20 | -580.95 |
| | | | ET+ | - | -599.94 | -598.65 | -596.78 | -601.57 | -598.04 | -596.83 | -597.05 | -597.23 | -599.05 | -596.82 | -524.39 | -596.82 |
| **VVS** | -736.73 | -735.22 | - | -767.83 | -730.03 | -754.29 | -731.47 | -746.46 | -755.80 | -730.06 | -732.99 | -735.27 | -732.87 | -744.20 | -641.64 | -737.98 |
| | | | ET+ | - | -760.75 | -764.12 | -761.27 | -764.98 | -762.15 | -760.58 | -761.90 | -761.75 | -761.95 | -760.89 | -669.51 | -760.89 |
| **Amy** | -2239.68 | -2372.46 | - | -2322.09 | -2317.26 | -2314.00 | -2276.71 | -2323.08 | -2330.33 | -2238.72 | -2238.99 | -2253.95 | -2280.78 | -2273.06 | -2028.50 | -2247.58 |
| | | | TV+ | -2337.51 | -2388.73 | -2330.90 | -2325.83 | -2348.95 | - | -2324.69 | -2324.55 | -2331.39 | -2360.47 | -2325.17 | -2112.81 | -2326.49 |
| | | | TV+nST+ | -2400.95 | - | -2381.51 | -2385.18 | -2385.18 | - | -2382.65 | -2382.53 | -2387.46 | -2381.51 | -2381.50 | -2150.11 | -2395.95 |
| | | | TV+AT+ET+ | - | - | -2393.94 | -2397.44 | -2397.44 | - | -2393.74 | -2393.72 | -2393.90 | -2393.94 | -2396.33 | -2161.91 | -2396.33 |
| **vmPFC** | -2292.75 | -2366.00 | - | -2340.14 | -2348.15 | -2327.19 | -2309.24 | -2354.44 | -2345.23 | -2286.66 | -2293.21 | -2297.90 | -2312.54 | -2305.47 | -2108.47 | -2298.28 |
| | | | AP+ | -2370.85 | -2371.35 | -2355.95 | -2389.60 | - | -2362.61 | -2358.72 | -2358.69 | -2347.65 | -2354.81 | -2350.36 | -2149.25 | -2358.10 |
| | | | AP+AB+ | -2387.28 | -2384.78 | -2384.56 | - | - | -2384.78 | -2383.20 | -2387.06 | -2388.71 | -2382.39 | -2383.69 | -2177.70 | -2391.48 |
| **ACC** | -2050.26 | -2292.87 | - | -2123.23 | -2213.52 | -2132.29 | -2071.41 | -2224.68 | -2137.06 | -2043.06 | -2060.56 | -2059.25 | -2131.92 | -2086.43 | -1893.75 | -2052.87 |
| | | | AP+ | -2240.72 | -2289.11 | -2231.99 | -2284.46 | - | -2227.37 | -2233.28 | -2253.11 | -2217.89 | -2250.82 | -2222.83 | -2036.43 | -2224.99 |
| | | | AP+nST+ | -2317.67 | - | -2288.41 | -2299.75 | - | -2299.75 | -2289.31 | -2292.42 | -2281.94 | -2281.96 | -2283.94 | -2087.96 | -2305.39 |
| | | | AP+nST+ET+ | - | - | -2310.66 | -2310.88 | - | -2310.88 | -2311.97 | -2314.83 | -2310.52 | -2311.50 | -2313.75 | -2115.92 | -2313.75 |
| **aIns** | -1387.27 | -1569.49 | - | -1435.42 | -1496.30 | -1434.02 | -1387.97 | -1540.58 | -1441.83 | -1380.11 | -1394.42 | -1390.16 | -1435.53 | -1408.01 | -1261.73 | -1387.94 |
| | | | AP+ | -1545.03 | -1572.27 | -1536.46 | -1560.92 | - | -1535.32 | -1543.32 | -1567.10 | -1535.10 | -1548.12 | -1534.66 | -1374.58 | -1539.18 |
| | | | AP+nST+ | -1582.81 | - | -1565.64 | -1569.63 | - | -1569.63 | -1569.94 | -1580.08 | -1565.43 | -1565.74 | -1565.25 | -1393.49 | -1580.58 |
| | | | AP+nST+ET+ | - | - | -1577.88 | -1575.91 | - | -1575.91 | -1576.93 | -1585.17 | -1576.70 | -1577.40 | -1579.21 | -1404.64 | -1579.21 |
| | | | AP+nST+ET+ | - | - | -1578.81 | -1577.93 | - | -1577.93 | -1578.11 | - | -1578.11 | -1578.97 | -1578.93 | -1405.69 | -1578.93 |
| **pIns** | -1741.87 | -2319.59 | - | -1841.54 | -2183.59 | -1919.5 | -1821.63 | -2034.84 | -1884.29 | -1737.7 | -1755.68 | -1768.59 | -1983.73 | -1830.78 | -1672.32 | -1736.80 |
| | | | nST | -2294.65 | - | -2247.15 | -2177.40 | -2289.92 | -2296.49 | -2178.27 | -2198.34 | -2207.35 | -2176.96 | -2212.04 | -2018.71 | -2212.67 |
| | | | nST+ET | - | - | -2295.48 | -2297.78 | -2342.33 | -2311.86 | -2287.41 | -2300.14 | -2299.12 | -2287.55 | -2287.41 | -2119.58 | -2287.41 |
| | | | nST+ET+AP | - | - | -2306.10 | -2323.47 | - | -2323.47 | -2307.27 | -2297.62 | -2286.37 | -2283.35 | -2293.74 | -2106.30 | -2305.95 |

Provided are the most likely combination of pattern component models (IPCs) to explain observed representational patterns for each ROI. Model fitting was performed by identifying the best individually fitting IPC, then iteratively adding remaining IPCs. IPCs held from prior levels of analyses are indicated in under 'Included+' (4th column from left), with the BIC score resulting from combining the held IPCs independently with all remaining IPCs displayed in the right columns. Significantly improved fit due to the addition of an additional IPC is indicated by ΔBIC > 2. Note, identified final IPC combinations predicted the observed data significantly better than a combination of all IPCs ('All'), and a single uniform predictor ('None').

**Supplementary Table ST3:** Alternate GBFS-BIC Paths: S1 & pIns

| | | | | **Information Pattern Component** – BIC Score | | | | | | | | | | | | |
|---|---|---|---|---|---|---|---|---|---|---|---|---|---|---|---|---|
| **ROI** | **None** | **All** | **Included+** | **ET** | **nST** | **ST** | **AC** | **AP** | **TV** | **PE** | **NE** | **AV** | **Sa** | **FS** | **VE** | **TA** |
| **S1** | -1034.08 | -1389.76 | - | -1119.00 | -1319.74 | -1195.3 | -1120.78 | -1188.25 | -1142.20 | -1036.30 | -1038.533 | -1062.32 | -1231.40 | -1128.35 | -952.89 | -1027.03 |
| | | | AP+ | -1405.82 | a | -1386.70 | -1322.49 | -1356.94 | -1398.61 | -1320.73 | -1323.06 | -1344.03 | -1319.99 | -1359.95 | -1198.08 | -1327.22 |
| | | | AP+nST+ | b | a | -1413.71 | -1398.70 | -1410.78 | -1412.63 | -1401.79 | -1403.37 | -1412.35 | -1401.70 | -1402.46 | -1269.41 | -1402.46 |
| | | | AP+nST+TV+ | b | a | -1410.12 | -1409.90 | -1409.90 | c | -1405.47 | -1405.93 | -1407.24 | -1410.12 | -1406.74 | -1275.21 | -1406.74 |
| **pIns** | -1741.87 | -2319.59 | - | -1841.54 | -2183.59 | -1919.50 | -1821.63 | -2034.84 | -1884.29 | -1737.77 | -1755.68 | -1768.59 | -1983.73 | -1830.78 | -1672.32 | -1736.80 |
| | | | nST+ | -2294.65 | a | -2247.15 | -2177.40 | -2289.92 | -2296.49 | -2178.27 | -2198.34 | -2207.35 | -2176.96 | -2212.04 | -2018.71 | -2212.67 |
| | | | nST+TV+ | -2311.86 | a | -2289.91 | -2323.47 | -2323.47 | b | -2298.02 | -2289.78 | -2295.92 | -2289.91 | -2289.50 | -2118.30 | -2301.72 |
| | | | nST+TV+AB+ | -2339.41 | a | -2316.92 | c | -2323.47 | b | -2317.21 | -2331.62 | -2323.07 | -2316.92 | -2316.49 | -2137.58 | -2329.016 |
| | | | nST+TV+AB+ET+ | d | a | -2332.17 | c | -2339.41 | b | -2337.65 | -2339.91 | -2332.34 | -2332.17 | -2333.32 | -2152.38 | -2333.32 |
| | | | - | -1841.54 | -2183.59 | -1919.50 | -1821.63 | -2034.84 | -1884.29 | -1737.77 | -1755.68 | -1768.59 | -1983.73 | -1830.78 | -1672.32 | -1736.80 |
| | | | nST+ | -2294.65 | a | -2247.15 | -2177.40 | -2289.92 | -2296.49 | -2178.27 | -2198.34 | -2207.35 | -2176.96 | -2212.04 | -2018.71 | -2212.67 |
| | | | nST+TV+ | -2311.86 | a | -2289.91 | -2323.47 | -2323.47 | b | -2298.02 | -2289.78 | -2295.92 | -2289.91 | -2289.50 | -2118.30 | -2301.72 |
| | | | nST+TV+AP+ | -2339.41 | a | -2316.92 | -2323.47 | c | b | -2317.21 | -2331.62 | -2323.07 | -2316.92 | -2316.49 | -2137.58 | -2329.01 |
| | | | nST+TV+AP+ET+ | d | a | -2304.62 | -2339.41 | c | b | -2308.88 | -2307.63 | -2304.79 | -2304.62 | -2305.74 | -2132.64 | -2305.74 |

With no clear IPC providing optimum fit at the 2nd and 3rd level of GBFS-BIC analyses (i.e. $IPC_1 - IPC_2 < 2$), search paths for each statistically equivalent option were completed. The IPC combination providing the lowest BIC at completion of the path was defined as the best fitting combination.

**Supplementary Table ST4:** *IPC* coefficient weights*: Full sample*

| | Regression | | Information Pattern Component weighting – β Coefficients | | | | | | | | | | | | | | |
|---|---|---|---|---|---|---|---|---|---|---|---|---|---|---|---|---|---|
| **ROI** | **$R^2$** | **P value** | **ET** | **nST** | **ST** | **AC** | **AP** | **TV** | **PE** | **NE** | **AV** | **S** | **FS** | **VE** | **TA** | **lST** | **rST** |
| **S1** | .247 | 2.8e-50 | 0.056 | 0.128 | 0.039 | - | - | - | - | - | - | - | - | - | - | n/a | n/a |
| **lS1** | .237 | .00053 | 0.051 | 0.136 | - | - | 0.045 | - | - | - | - | - | - | - | - | n/a | - |
| **rS1** | .240 | 7.3e-7 | 0.067 | 0.131 | - | - | - | - | - | - | - | - | - | - | - | 0.050 | n/a |
| **S2** | .403 | 9.1e-88 | 0.040 | 0.183 | - | - | 0.101 | - | -0.030 | - | - | - | - | - | - | n/a | n/a |
| **V1** | .021 | 2.0e-8 | 0.059 | - | - | - | - | - | - | - | - | - | - | - | - | n/a | n/a |
| **VVS** | .026 | 6.1e-10 | 0.061 | - | - | - | - | - | - | - | - | - | - | - | - | n/a | n/a |
| **Amy** | .120 | .00036 | 0.033 | 0.046 | - | - | - | 0.020 | - | - | - | - | - | - | - | n/a | n/a |
| **vmPFC** | .075 | 8.9e-21 | - | - | - | 0.052 | 0.075 | - | - | - | - | - | - | - | - | n/a | n/a |
| **ACC** | .184 | 1.1e-10 | 0.037 | 0.058 | - | - | 0.062 | - | - | - | - | - | - | - | - | n/a | n/a |
| **aIns** | .147 | 8.2e-13 | 0.028 | 0.044 | - | - | 0.126 | - | - | -0.036 | - | - | - | - | - | n/a | n/a |
| **pIns** | .356 | 6.0e-66 | 0.047 | 0.112 | - | - | 0.070 | - | - | - | - | - | - | - | - | n/a | n/a |
